# Reliable variant calling during runtime of Illumina sequencing

**DOI:** 10.1101/387662

**Authors:** Tobias P. Loka, Simon H. Tausch, Bernhard Y. Renard

**Author notes:** Correspondence to Bernhard Y. Renard, Robert Koch Institute, Nordufer 20, 13353 Berlin. German Federal Institute for Risk Assessment (BfR), Department of Biological Safety, Berlin, Germany (Simon H. Tausch).

## Abstract

The sequential paradigm of data acquisition and analysis in next-generation sequencing leads to high turnaround times for the generation of interpretable results. We combined a novel real-time read mapping algorithm with fast variant calling to obtain reliable variant calls still during the sequencing process. Thereby, our new algorithm allows for accurate read mapping results for intermediate cycles and supports large reference genomes such as the complete human reference. This enables the combination of real-time read mapping results with complex follow-up analysis. In this study, we showed the accuracy and scalability of our approach by applying real-time read mapping and variant calling to seven publicly available human whole exome sequencing datasets. Thereby, up to 89% of all detected SNPs were already identified after 40 sequencing cycles while showing similar precision as at the end of sequencing. Final results showed similar accuracy to those of conventional *post-hoc* analysis methods. When compared to standard routines, our live approach enables considerably faster interventions in clinical applications and infectious disease outbreaks. Besides variant calling, our approach can be adapted for a plethora of other mapping-based analyses.

## Introduction

Common workflows for the analysis of Illumina next-generation sequencing (NGS) data can only be applied after sequencing has finished. Besides the time needed for sample preparation, this sequential paradigm of data acquisition and analysis is one of the main bottlenecks leading to high turnaround times. For time-critical applications, it is crucial to massively reduce the time span from sample receipt to interpretable analysis results. Examples for such time-critical analyses range from the differential diagnosis of genetic disorders in infants^1, 2, 3, 4^, to the determination of *M. tuberculosis* drug resistances^5^, and to the identification of pathogens, virulence factors, drug resistances and paths of disease transmission in infectious disease outbreaks^6, 7^. While having considerably higher turnaround times than targeted approaches such as molecular tests, NGS provides a more open view as well as more extensive and reliable results. During bioinformatics analysis of NGS data, read mapping and variant calling are crucial steps to obtain genetic information that is essential for the treatment of a patient, including strain level classification and drug resistances of a pathogen or the presence of genetic disorders that are known to be associated with specific disease characteristics.

While HiLive^8^, the predecessor of our new algorithm HiLive2, delivered results at the end of sequencing, HiLive2 can produce read mapping output for arbitrary sequencing cycles while still sequencing. At the same time, the new algorithm is faster, more accurate and enables scalability to large reference genomes such as the complete human reference. The recently published software LiveKraken^9^ already gives k-mer based taxonomic classification results for arbitrary sequencing cycles. However, while LiveKraken provides valuable information about the microbial composition of a sample, the results do not allow for complex reference-based follow-up analyses such as variant calling or the analysis of drug resistances. Alternative approaches to obtain read mapping results for Illumina data while still sequencing, such as rapid pulsed whole genome sequencing^4^, lack sufficient scalability for high amounts of data and large reference genomes and are therefore only suitable for special use cases. At the same time, the incremental approach of HiLive2 provides higher flexibility in the choice of output cycles which can even be modified during the runtime of the sequencer. The use of specialized hardware, such as field programmable gate arrays (FPGAs) that are for example used in the DRAGEN system^3^ could generally overcome the lack of scalability and speed for intermediate analyses but come with additional costs, either for purchase and infrastructure of local solutions or for the use of a cloud system. At the same time, such approaches are usually not algorithmically optimized for analyzing incomplete data. Additionally, cloud solutions as provided by DRAGEN can be problematic with regard to data protection guidelines in many countries.

A different tool, TotalReCaller^10^, implements a similar algorithmic idea as HiLive2. TotalReCaller uses an FM-index based alignment approach to perform reference-based base calling. While TotalReCaller’s alignment approach only allowed for substitutions, its successor Gappy TotalReCaller^11^ also considers insertions and deletions. However, as TotalReCaller’s main objective is to improve base calling it does not produce actual alignment output. At the same time, the authors describe the need for an FPGA-implementation to make it suitable for real-time applications.

When compared to Oxford Nanopore sequencing Technology (ONT), which enables real-time analysis by design, Illumina sequencing provides higher scalability at lower costs and with much lower error rates. Therefore, while being a promising technology for real-time analysis in the future, sequencing technology, protocols and computational analysis for ONT need to be further established and improved to become a viable alternative for many scenarios.

The workflow described in this study is based on real-time read mapping results with our novel algorithm HiLive2 followed by fast and accurate variant calling with xAtlas^12^. Thereby, live results can be obtained several hours before all data are written by the sequencer and provide increasing insights into the sample over sequencing time. This study describes the application of our workflow to human whole exome sequencing data, showing the scalability of our approach for high amounts of complex data and large reference genomes. However, our approach can generally be adapted for different types of sequencing methods such as whole genome sequencing (WGS) or amplicon sequencing and a plethora of different mapping-based analysis methods. While application to arbitrary sample types and reference genomes is possible in general, the latency of real-time results strongly depends on the size and complexity of the used reference genome, the number of reads pre tile, the parameter settings made and the available hardware. Therefore, when planning real-time sequencing, it should be examined if the desired live analysis of a sample is possible under the given conditions.

Our new real-time read mapping software HiLive2 is publicly available under BSD-3-clause on https://gitlab.com/rki_bioinformatics/hilive2 and on Bioconda for easy installation (conda install –c bioconda hilive2).

## Results

### Implementation and experimental setup

To allow for faster NGS-based diagnosis and treatment, we developed a new real-time read mapping algorithm that generates high-quality results. We combined our new software with a fast variant caller to produce high-quality variant calls based on intermediate read mapping results, while sequencing is still running. This approach allows reliable and fast variant calling results without reducing the final sequencing coverage or quality. Therefore, we adapted our real-time read mapper HiLive^8^ that gives output at the end of sequencing, using a novel algorithm based on the efficient FM-index^13^ implementation of the SeqAn library^14^ for continuously analyzing sequencing results during runtime. The new version (HiLive2) achieves scalability to larger indices such as the complete human reference genome. At the same time, the algorithm comes with improved performance in terms of runtime, memory and data storage and overcomes heuristic elements that were present in previous version of HiLive. The high scalability and accuracy of HiLive2 enable the combination of real-time read mapping results with complex follow-up analyses that have not been possible with the previous version. To demonstrate the power of such analyses, we performed variant calling on seven whole exome sequencing data sets of the human individual NA12878 from the CEPH Utah Reference Collection (cf. Table 1) using the real-time read mapping results of HiLive2 as input data.

**Table 1.**
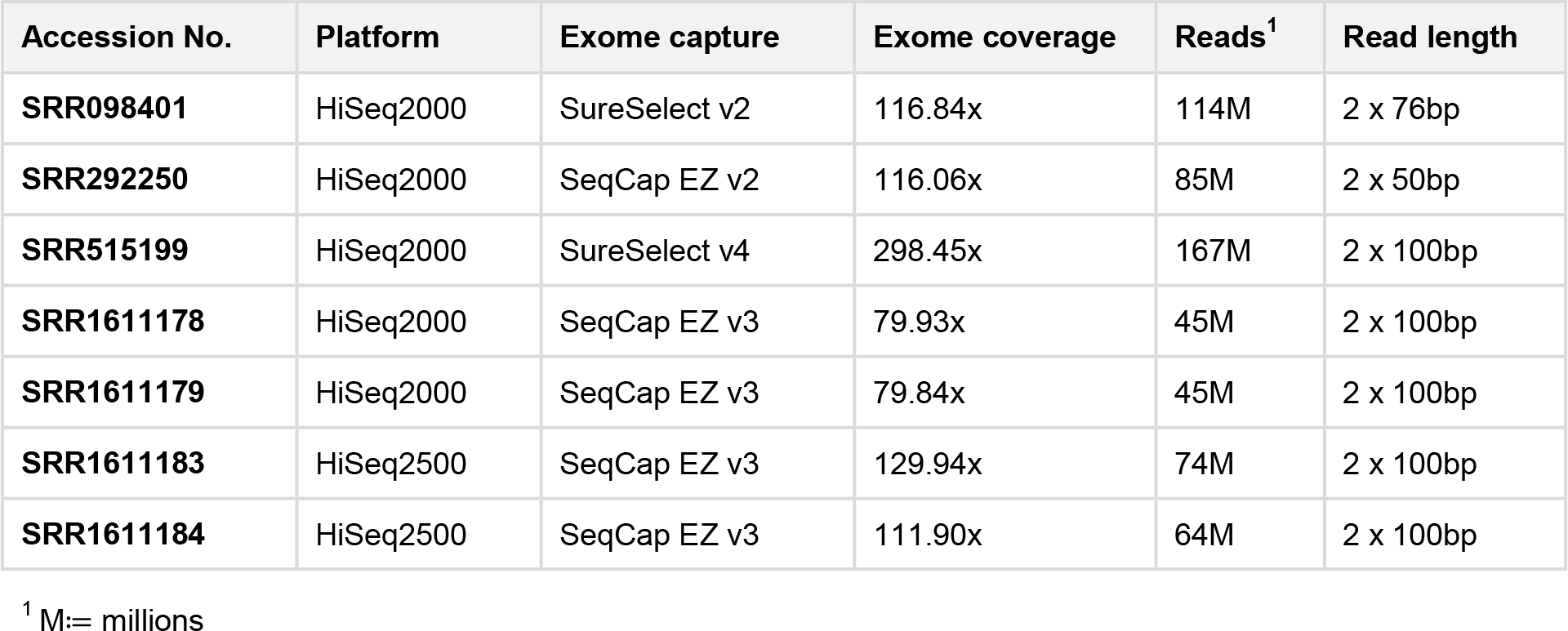
Summary of data sets evaluated in this study. Information about sequencing platform, exome capture and coverage were adopted from Hwang et al. (2015)^15^.

For variant calling, we used the fast variant caller xAtlas which shows comparable accuracy to established methods at much lower runtime^12^. We compared our results to read mapping with Bowtie 2^16^ and variant calling with either xAtlas or GATK 3.3.0^17^ for the same data sets. For Bowtie 2 + GATK, we took the results from Hwang et al. (2015)^15^ following the GATK best practice procedure using Picard ReorderSam (http://broadinstitute.github.io/picard/) and GATK IndelRealigner, BaseRecalibrator and HaplotypeCaller. Accuracy was determined by comparing the results to the well-established high-confident variant calls for the human individual NA12878 published by the Genome in a Bottle (GIAB) consortium^18^. As benchmarking method we used the area under the precision-recall curve (APR).

### Accuracy of real-time results

In Illumina sequencing, all reads are sequenced in parallel. In each so-called sequencing cycle, sequence information of one additional nucleotide is obtained for all reads. Thus, the current length of a read equals the number of the respective cycle (e.g., 40 nucleotides after cycle 40). To demonstrate the capability of our approach to provide interpretable results during runtime, we applied our workflow at different stages of sequencing. We expected our live results to show higher accuracy for higher cycles due to the increasing amount of available sequence information. At the same time, we analyzed whether the detected variants in early sequencing cycles are as reliable as variants called at the end of sequencing. This is a crucial criterion for the proposed workflow since interpretation of live results is only meaningful when based on reliable variant calls. Therefore, besides comparing the APR values of different sequencing cycles, we also examined precision and recall separately.

Fig. 1a shows the progression of the APR values for SNP calling in all analyzed data sets with increasing sequencing time. In cycle 30, sequence information was not sufficient to call any variants with the given parameter settings for six of seven data sets. For data set SRR292250, read mapping parameters were adapted by HiLive2 automatically due to the short read length of 50bp. This led to earlier results after 30 cycles, while first results were available after cycle 40 for all other data sets. Results show a continuous increase of the APR values for all cycles of the first read. In cycle 75, an APR larger than 0.9 was achieved for all data sets with sufficient read length. Afterwards, the APR values continue increasing moderately. When regarding the progression of precision (Fig. 1b) and recall (Fig. 1c) over sequencing time separately, it can also be also observed that lower APR values for earlier sequencing cycles are mainly caused by a lower recall while precision changes only slightly with more sequence information available. The same conclusions are supported by the individual precision-recall curves for all data sets which show a large increase of the recall but only minor changes of specificity over sequencing time (cf. Supplementary Fig. 1). This indicates that live results are highly reliable and can therefore serve for early interpretation and problem-specific follow-up analyses.

**Fig. 1:**
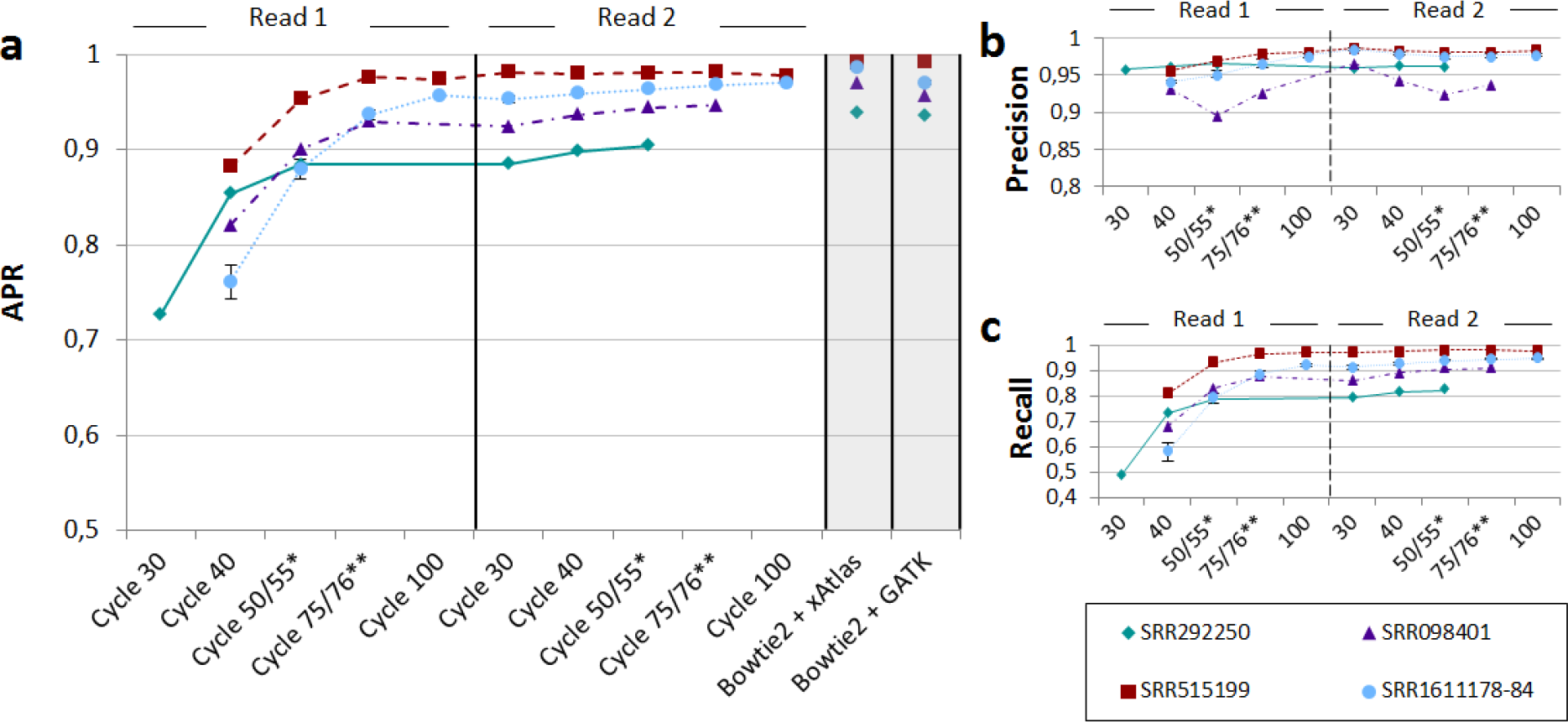
Area under a precision-recall curve (APR) for SNP calling in seven data sets at different sequencing cycles. SNP calling was performed with xAtlas using real-time read mapping results of HiLive2. Results for the samples SRR1611178, SRR1611179, SRR1611183 and SRR1611184 were combined to a single data series due to their high similarity (SRR1611178-84). Error bars for this data series show the standard deviation. Reads of SRR292250 and SRR098401 were shorter than 2×100bp which leads to missing data points. The vertical, ticked line in the middle of the plot divides the first and second read. **a** The gray columns show APR values using Bowtie 2 for read mapping and xAtlas (left) and GATK (right) for variant calling. The data for Bowtie 2 + GATK were taken from Hwang et al. (2015)^15^. The real-time workflow with HiLive2 and xAtlas provides first results after 40 sequencing cycles (30 cycles for SRR292250). An APR greater than 0.9 is reached after 75 cycles for all data sets with a minimal read length of 75bp. Until end of sequencing, there is a moderate increase of the APR. **b** Precision with a quality threshold of 1 for variant calling with xAtlas. The results show no precision lower than 0.89 for all sequencing cycles. In general, precision increases only slightly over time. This indicates that results in early sequencing cycles are already reliable. **c** Recall with a quality threshold of 1 for variant calling with xAtlas. The results show strong improvements from the first results available until the end of the first read. The progression of all curves is similar to that of the APR curve (cf. Fig. 1a), indicating the correlation between those two measures. *Cycle 50 for SRR292250, cycle 55 for all other data sets. **Cycle 76 for SRR098401, cycle 75 for all other data sets.

The increasing number of SNP calls in subsequent cycles provides additional information for complementing the previous interpretation of the data. However, the final results with HiLive2 show slightly lower maximum recall values than the same workflow applied to read mapping results of Bowtie 2 (cf. Supplementary Fig. 1). This can be explained by the read mapping approach of HiLive2 which tolerates only a specified number of errors for a read. Thus, regions with a high number of variations may be lowly covered which leads to undetected variants. The same effect is somewhat stronger for indels as the mapping algorithm only tolerates indels with a maximum length of three nucleotides by default due to computational costs. While this behavior led to a lower recall than based on read mapping with Bowtie 2, the results showed comparable or higher precision (cf. Supplementary Fig. 1). Thus, although focussing on SNPs in this study, our workflow can also provide valuable insights about small indels.

### Turnaround time of the workflow

Besides the accuracy of results, turnaround time is the second crucial factor for NGS-based real-time analyses. Thereby, live results should be available as soon as possible after the data of the respective sequencing cycle was written without showing significant delay in any stage of sequencing.

We measured the turnaround time of real-time mapping with HiLive2 and subsequent variant calling with xAtlas for the same runs that delivered the accuracy results shown before. All computations were run on a 128-core machine (Intel® Xeon® CPU E5-4667 v4 @ 2.20 GHz, 45 M Cache) with 500GB RAM, using a maximum of 65 threads per data set. Fig. 2 shows an overview for the turnaround time of our workflow for different sequencing cycles for data sets SRR1611184 and SRR515199. For SRR1611184, variant calling results were available after a maximum of 35 minutes for all output cycles of the first read and a maximum of 52 minutes for all output cycles of the second read. With approximately 1 million reads per tile (or thread), these are realistic numbers for a real-case scenario using benchtop sequencing devices. For SRR515199, the latency is way higher reaching a maximum of more than three hours (193 minutes) for cycle 175. The higher latency originates from analyzing approximately 2.5 times as many reads per tile (or thread) as for data set SRR1611184. This shows that the latency of real-time results strongly depends on the number of reads that are analyzed per thread which varies with the available hardware and the used sequencing device. In general, the analysis of high coverage data sets can be significantly reduced by adapting the alignment parameters at the cost of accuracy. However, for the sake of comparability, we ran all data sets with the same parameter settings in this study leading to a higher latency for higher coverage data sets. Results for the other data sets are shown in Supplementary Fig. 2. Thereby, five of the seven data sets showed a maximum latency of less than one hour from data output to interpretable results for each output cycle.

**Fig. 2:**
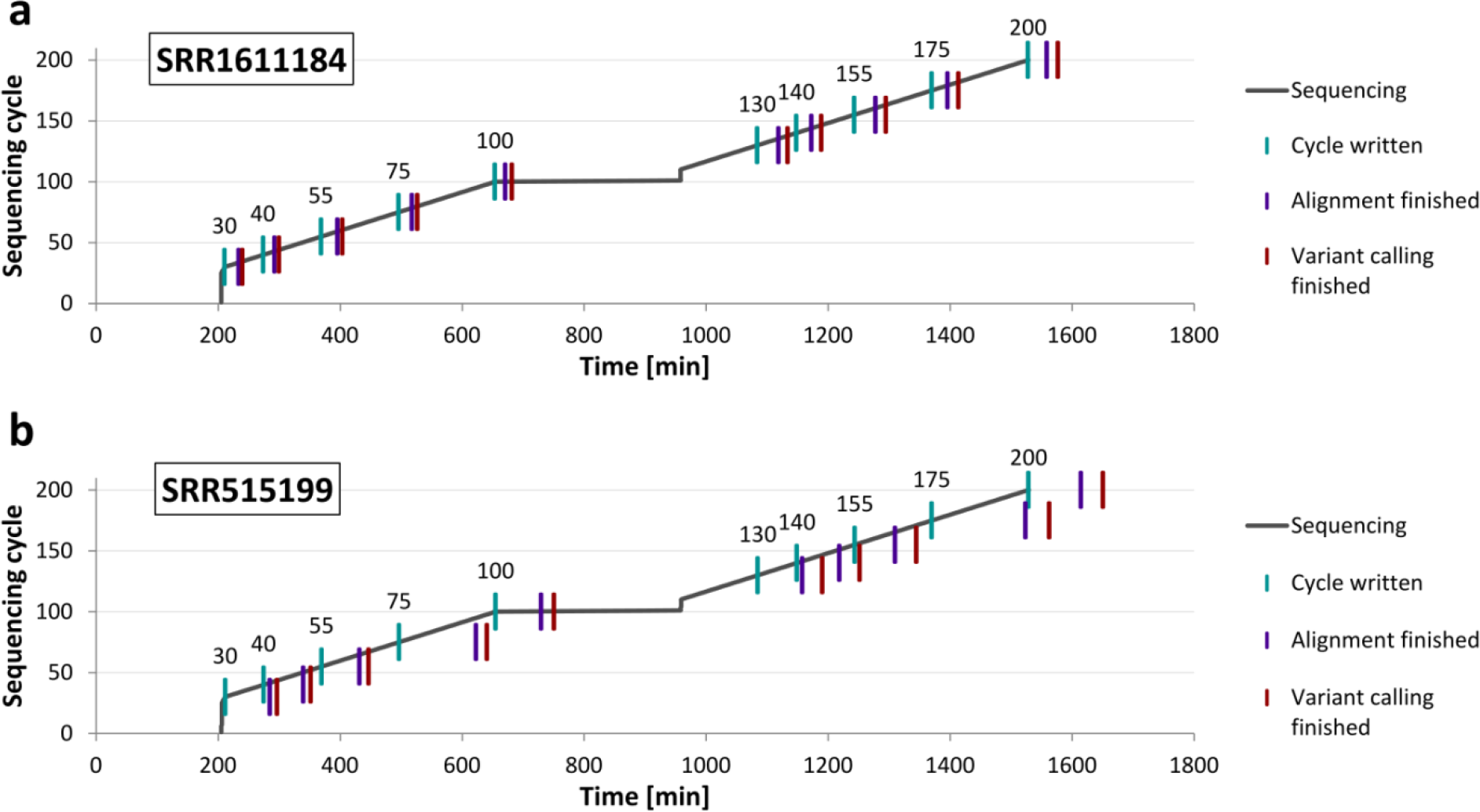
Turnaround time of our workflow for data sets SRR1611184 (a) and SRR515199 (b). For each cycle, the first vertical line indicates the time point when the data for the respective cycle was completely written. The second line shows when the alignment output of HiLive2 is written. The third line indicates the end of our workflow resulting in the output of variant calls for the respective cycle. Vertical lines with the same vertical position belong to the same output cycle.

## Discussion

In clinical applications and infectious disease outbreaks, the turnaround time of analyses is a critical factor for an effective treatment of patients. At the same time, a high analysis depth and an open perspective for unexpected findings are further crucial criteria in such scenarios. Therefore, despite its significantly higher turnaround times than alternative methods, NGS presents an established analysis method in several time-critical applications. For example, a comprehensive report of vancomycin resistant *Enterococcus faecium* infections in three patients was created in 48.5 hours including over-night culturing using an Illumina MiSeq benchtop sequencer^19^. Also in the field of acutely ill infants with suspected genetic diseases, there were impressive improvements in the applicability of NGS-based diagnostics including a 26 hours protocol for provisional molecular diagnosis^3^. However, even with a significant speed-up of the computational analysis using faster software or specialized hardware such as field-programmable gate arrays (FPGAs), a decrease of turnaround time is strictly limited due to the sequential paradigm of data creation and analysis. Motivated by this, Miller et al. (2015) introduced the idea to combine bioinformatics analysis using FPGAs, in particular the DRAGEN system, with the concept of rapid pulsed whole genome sequencing^4^ to achieve near real-time analysis results^3^. Such an approach would require a conversion of sequencing data to FASTQ or BAM/CRAM format for each desired output cycle as these are the only file formats supported by the DRAGEN system. At the same time, specialized hardware is required and licenses must be purchased. Alternatively, a cloud service can be used which can be problematic due to data protection guidelines and the required speed of data transfer. However, to the best of our knowledge there is no study describing a proof-of-principle for this idea. Another commercially available approach to speed up NGS analysis is implemented in the Sentieon Genomics Tools, an optimized version of BWA and GATK that overcomes the need of specialized hardware or cloud access^20^. On the other hand, this software is not specifically designed to produce real-time results and involves costs for licenses. Furthermore, both commercial systems are developed with a strong focus on variant calling for human samples. In contrast, the workflow presented in this study is open source, runs on a standard linux machine and allows for easy and flexible adaption of the workflow for different scenarios. At the same time it is highly scalable and provides high-quality analysis results without the need of acquiring specialized hardware, cloud access and licenses.

The results of our study demonstrate the enormous potential of our approach to reduce the turnaround time from sample arrival to meaningful analysis output by several hours up to days depending on the used sequencing device. Thereby, live results in very early stages of sequencing can already deliver highly confident results while the quantity of analysis results (i.e. the number of called variants in this study) increases with a growing number of sequenced nucleotides per read. Live analyses can therefore provide first relevant insights into the data while the analysis becomes more comprehensive with ongoing sequencing. The sensitivity of results in early sequencing cycles is thereby implicitly limited by the lower total coverage when compared to the full read length as well as the relatively high number of unambiguously mapped reads, especially in low complexity regions. To demonstrate the power of our approach, we showed its application to human whole exome sequencing data including real-time alignment of all reads to the full human reference genome hg19. We chose this type of data due to its complexity and computational demands as well as the availability of high-quality and extensively studied gold-standard data sets provided by the GIAB consortium. However, our approach is not restricted to human whole exome sequencing data and the presented use case of variant calling. We rather see an enormous potential of real-time read mapping to provide means for a wide range of complex follow-up analyses for various types of data. Still, due to current technical limitations of HiLive2 and runtime limitations of tools for subsequent analysis such as variant calling, certain types of analyses that require huge amounts of data are only feasible with limitations. For example, when applying the same SNP-calling workflow as shown for the WES data to a 30× WGS data set (SRR6808334)^21^ of the human individual NA12878 (cf. Supplementary Fig. 3), the latency of real-time results was up to six hours while achieving a maximum recall of 87% with 97% precision. Even for such analyses, valuable insights could be gained for early sequencing cycles (e.g., 40% recall with 90% precision after 55 cycles and 62% recall with 91% precision after 75 cycles), but given the high latency and comparatively low accuracy we only recommend our workflow for such data sets in exceptional situations and when no other options are available. However, future improvements, such as a decrease of I/O operations, in-tile multithreading and acceleration of alignment output, could significantly reduce the latency of HiLive2 and make it more applicable for higher throughput applications.

A second limitation of our approach comes up when dealing with multiplexed sequencing data which is usually applied for high-throughput applications to sequence more than one sample within one sequencing run. While HiLive2 provides demultiplexing functionality and can produce separate alignment files for each sample, the sequencing approach of Illumina limits the early assignment of reads by sequencing the barcodes after completing the first read. In doing so, it is not possible to distinguish between different samples before sequencing of the first read has finished. A change in the sequencing order is not trivial as sequencing the barcodes first would negatively influence the initial clustering that is performed during the first couple of sequencing cycles and relies on a wide variety of sequences which is not given for the barcodes, in particular when only having a low number of samples. When single-end sequencing is sufficient, this can be overcome by performing asymmetric paired-end sequencing. This means, that the first read is only sequenced for several base pairs to enable proper clustering, followed by the barcodes and the complete second read. While resulting in some additional runtime at the beginning of sequencing, this enables an early assignability of reads to the samples. A different conceivable solution could be to include a random sequence followed by inline barcodes at the beginning of the first read.

Alternative approaches to NGS for diagnosis can also be highly valuable for different scenarios. Molecular approaches are usually highly reliable and provide answers to specific questions in a very short timeframe and at much lower costs. For example, the detection of 25 genetic mutations in *M. tuberculosis* that confer to drug resistances can be finished in approximately two hours with a variation of the molecular GeneXpert test^5^. Even when providing live results, such short turnaround times are currently not feasible with NGS-based approaches due to the required time for sample preparation and clustering. Another interesting technology for time-critical applications is the Oxford Nanopore sequencing Technology (ONT). It was shown that metagenomic detection of viral pathogens can be achieved in less than six hours^22^. While ONT shows a high portability and much faster sample preparation as additional benefits, this and other current long-read technologies are still expensive and limited by their comparatively low coverage and high error rates. It is therefore hard to reliably identify lowly abundant pathogens, genetic variants, parallel infections or the presence of viral quasispecies. Thus, especially when it comes to these or other questions going beyond the identification of highly abundant pathogens in time-critical applications, real-time analyses for Illumina sequencing can be of great benefit.

Concluding, we consider our new real-time workflow for Illumina sequencing to be a complementary method to molecular tests and ultra-portable, long-read sequencing for time-critical analyses. It fills the current gap of short turnaround times, an open-view perspective and high sequencing coverage which is essential for a plethora of applications such as pathogen identification and characterization, identification of acute genetic diseases or epidemiological analyses. Therefore, our approach is an important step for improving the ability for fast interventions in exceptional clinical situations, personalized medicine and infectious disease outbreaks.

## Methods

In this chapter, we provide a brief description of our workflow. More detailed information for each step as well as direct links to the data and the executed commands can be found in the supplementary material. All software versions are listed in the *Software versions* section.

### Implementation of HiLive2

HiLive2 is based on a novel algorithm based on the efficient FM-index implementation of the SeqAn library^14^. HiLive2 provides five different modes to select a focus on runtime (fast or very-fast), accuracy (accurate or very-accurate) or a trade-off of both (balanced). Parameter decisions are made automatically by HiLive2 based on the selected mode, the size of the reference genome and the read length. The influenced parameters include the length of an initial, error-free *k*-mer (anchor-length), the intervals of creating new seeds (seeding-interval), the minimum alignment score (min-as) and the intervals of increasing the number of errors (error-interval). However, all parameters can be refined or completely manually set by the user to enable even faster or more accurate computations.

HiLive2 follows the seed-and-extend alignment approach. The first seeds are created when sufficient bases are available for all reads to create an anchor of the specified length (initial seeding). Thereby, anchors are error-free matches of the most recently sequenced *k*-mer in the index. New alignment anchors of the same length are created in specified intervals (non-initial seeding). By default, the selected intervals produce overlapping anchors. In the following cycles after the creation of an anchor, the alignments are extended in the direction of sequencing only. This means that alignments originating from non-initial seeding have an unaligned region at the beginning of the reads which are reported as softclips. During extension, the minimum score of the alignment decreases with ongoing sequencing. This means that more errors are permitted for longer sequences. This approach leads to a massive reduction of the search space in early sequencing cycles while having only minor effects on the final results as the mapping positions of reads with too many errors at the beginning can still be determined based on a non-initial seeding step. However, the actual alignment information for the errorneous regions is lost in this case but can be covered by other reads having these regions being placed in the middle or at the end of the read.

HiLive2 is an all-mapper by design. This means that all alignments within the search space specified by the parameter setting can be found and reported. However, the default output option of HiLive2 is to report only one best alignment for each read. For most analyses, this output option is the expected behavior. Output is written in the well-established BAM or SAM format. For all output cycles, temporary files are stored and can be used to efficiently produce output with different options (e.g., writing all alignments instead of one best alignment for each read). This functionality is particularly useful for explorative analyses. Additionally, these temporary files can be used as an entry point to continue the alignment if the process crashed or had to be interrupted.

### Data download and conversion

The human reference genome hg19 was obtained from NCBI and only considered chromosomes 1-22, X and Y. Alternative regions were omitted. The sequences were stored in a single multi-FASTA file. For the evaluation of variant calls with RTG Tools^23^, the reference genome was converted to SDF format.

The sequencing data sets of the individual NA12878 were downloaded from EBI in FASTQ format. For read mapping with HiLive2, read pairs were converted to Illumina base call file format (BCL), distributed on one lane and 64 tiles. There were four different definitions for exome capture region definition required for the different data sets (cf. Table 1). The regions were obtained in BED format from the respective producer, if available. Whenever multiple definition files were provided, the primary target regions were selected.

Gold standard variants for the individual NA12878 were downloaded from the Genome in a Bottle (GIAB) consortium^18^ and regularized with the vcfallelicprimitives tool of VCFtools^24^. SNPs and indels of the gold standard were stored in two separated files and filtered out against the exome capture regions using Bedtools^25^ intersect. The resulting files were used as the gold standard for data sets using the respective exome capture definition. During the evaluation of the results, only variant calls in high confidence homozygous regions which were obtained from GIAB were considered.

### Real-time read mapping with HiLive2

The index of human reference genome hg19 for HiLive2 was built with default parameters. The creation of base call files by the sequencing machine was simulated using a script for sequencing simulation with a sequencing profile for HiSeq2500 machines in rapid mode and using dual barcodes. As no barcodes were present in our data sets, no data was written by the sequencing simulator for the respective cycles. HiLive2 was run in fast mode allowing faster turnaround times at the expense of slightly lower recall. Technical parameters as lanes, tiles and read length were set according to the data sets. In general, we chose cycles 30, 40, 55, 75 and 100 for each of the two reads as output cycles. For data sets with read lengths other than 2 × 100bp, we adapted the output cycle numbers to 30, 40 and 50 (SRR292250) or 30, 40, 55 and 76 (SRR098401). We used the recommended number of threads (1 thread per tile) for HiLive2 resulting in 64 threads for all data sets.

### Read mapping with Bowtie 2

The index of human reference genome hg19 for Bowtie 2 was built with default parameters. Read mapping with Bowtie 2 was performed with default parameters using 10 threads.

### Variant calling with xAtlas

Variant calling with xAtlas was performed for each chromosome individually. Therefore, the alignment files of HiLive2 or Bowtie 2 were split in 24 files (one for each chromosome). The resulting files were sorted and indexed using samtools^26^. Afterwards, variants were called with xAtlas for the respective exome capture regions using default parameters. Sorting, indexing and variant calling was performed with 24 threads (1 per chromosome). The resulting VCF files were merged using VCFLIB vcf-concat (https://github.com/vcflib/vcflib) for SNVs and indels separately.

### Measure of turnaround time

The sequencing simulation script provides timestamps for each written sequencing cycle. These timestamps were compared to the system time stamps for the last modification of the alignment output files of HiLive2. The time span between both time stamps describes the alignment delay of HiLive2. Additionally, we measured the clock time of the xAtlas pipeline. The sum of sequencing time until the respective cycle, the alignment delay of HiLive2 and the clock time of xAtlas yields the overall turnaround times of our workflow.

### Evaluation with RTG Tools

We used the vcfeval program of RTG Tools for the validation of variant calling results. We used the gold standard for the respective data set (depending on the used exome kit) as baseline and the variant calling output of xAtlas or GATK as call. The human reference hg19 in SDF format was used as reference template. Only variant calls being included in the high-confidence regions for individual NA12878 provided by the GIAB consortium were considered for validation. We ran RTG Tools with 24 threads and used the squash-ploidy and all-records parameters. For variant calls produced by xAtlas, we additionally defined QUAL as the field for variant call quality. For GATK, the GQ field is chosen by default. RTG Tools vcfeval returns a list of statistical measures for different thresholds of the variant call quality field, including precision and recall. These values served as input for the precision-recall curves shown in Supplementary Fig. 1 and used for the calculation of the area under the precision-recall curves (APR).

### Statistical Measures

We used precision and recall values for the validation of our approach. True positives (TP) describe the number of correctly detected variants. False negatives (FN) are the number of undetected variants. False positives (FP) are the number of variants that were detected by our pipeline but are not contained in the gold standard.

Recall is the relative number of variants of the gold standard that were found by our approach (TP / (TP + FN)). Precision is the fraction of variants called by our approach that are also present in the gold standard (TP / (TP + FP)).

### Software versions

Table 2 shows the software versions used for this study.

**Table 2.**
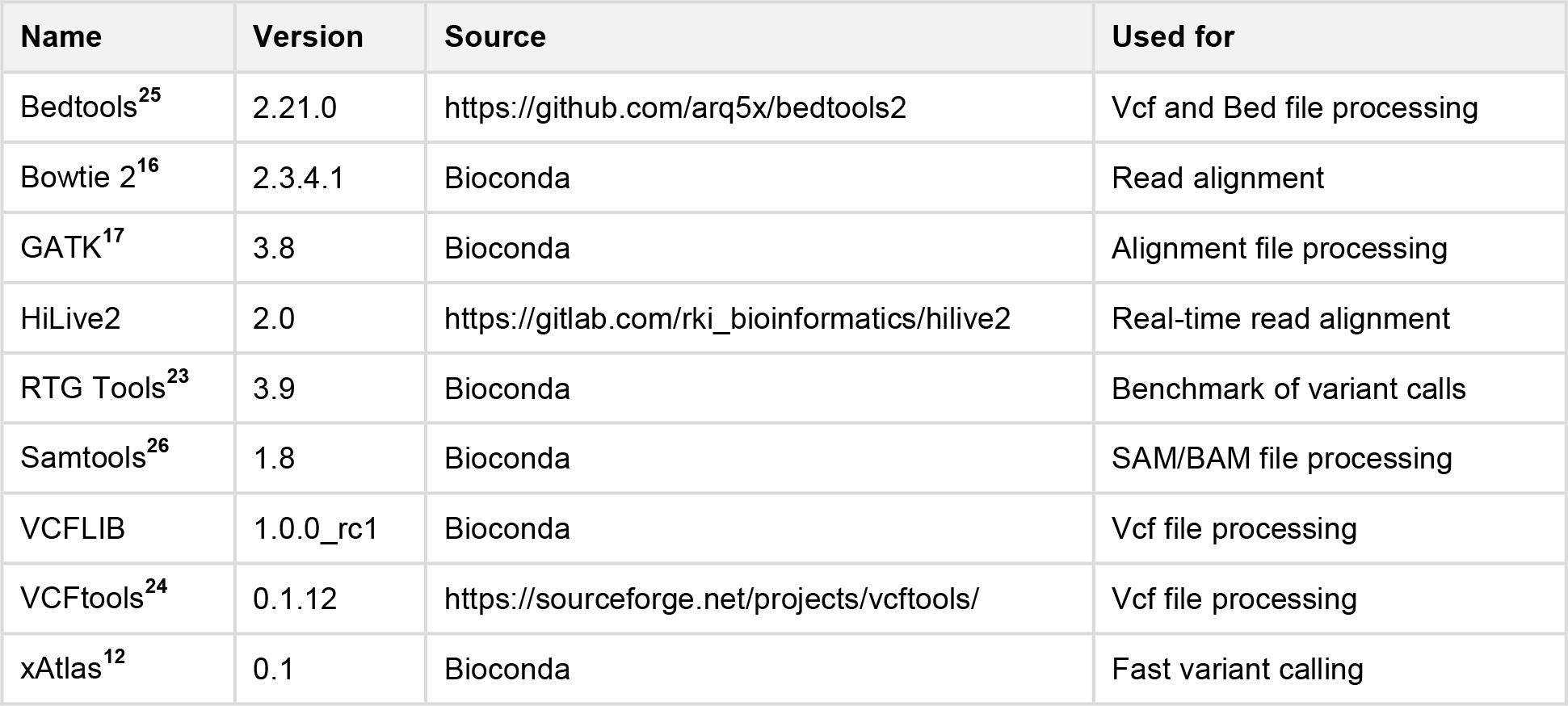
Table 2 List of software used in this study. Software with italic Bioconda was installed with the environment management software conda (https://conda.io) and obtained from the Bioconda channel^27^.

## Code availability

The source code of HiLive2 is available for public download on https://gitlab.com/rki_bioinformatics/hilive2 and comes with extensive documentation and sample data. HiLive2 is also available on BioConda for easy installation (conda install –c bioconda hilive2). Scripts that were used for our analyses are provided as Supplementary Software.

## Data availability

Sequencing data of the individual NA12878 is publicly available on the NCBI Short Read Archive and on the EBI FTP server. Gold standard variant calls are publicly available from the Genome in a Bottle Consortium. Human reference genome hg19 was obtained from the NCBI FTP server. Exome capture targets are available from the manufacturers or from third party resources.

## Acknowledgements

This work was supported by the German Federal Ministry of Health [IIA5-2512-FSB-725 to B.Y.R. and 2515NIK043 to S.H.T.] and the BMBF Computational Life Sciences program. We thank Andrea Thürmer and Aleksandar Radonić for their input concerning technical aspects of Illumina sequencing. We further thank Wojciech Dabrowski for infrastructural support and Thilo Muth for critical reading of the manuscript and his highly valuable suggestions. We also thank Martin Lindner, Jakob Schulze, Benjamin Strauch, and Kristina Kirsten for their work on HiLive.

## Contributions

B.Y.R. and T.P.L. conceived the study. T.P.L. performed the implementation of HiLive2. S.H.T. continuously supported the development of HiLive2 and evaluated the performance of HiLive2 on different types of data. T.P.L. designed the workflow and performed the analyses. All authors were involved in the preparation of the manuscript and approved the final version.

