## Supplemental Figures 1-3 and Supplemental Methods for "Reliable variant calling during runtime of Illumina sequencing"

**--- Supplementary Material ---**

#### Supplementary Fig. 1

##### SRR098401

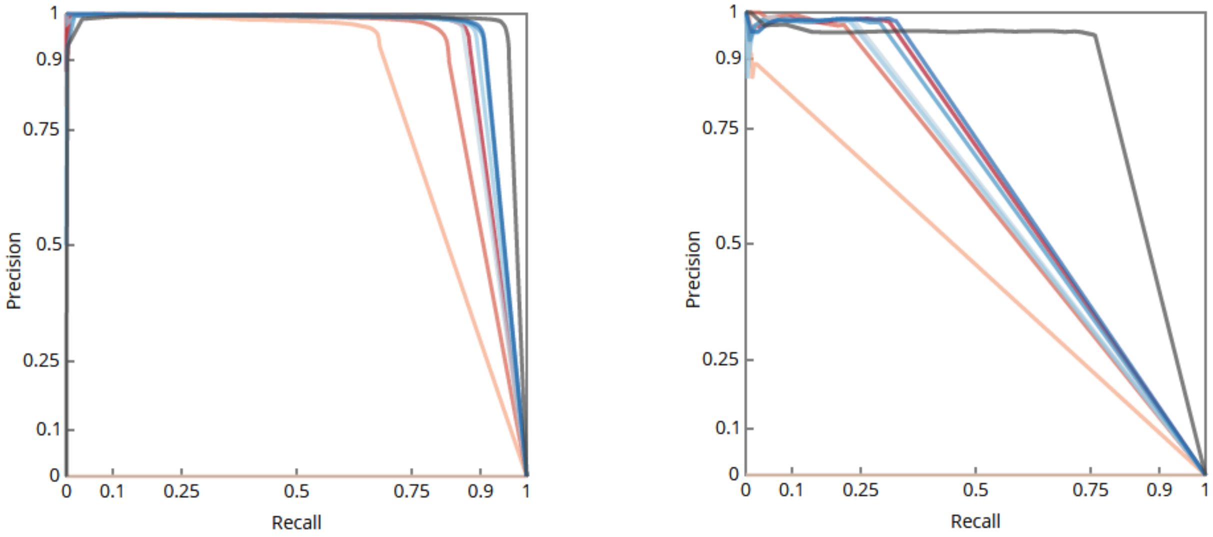

##### SRR292250

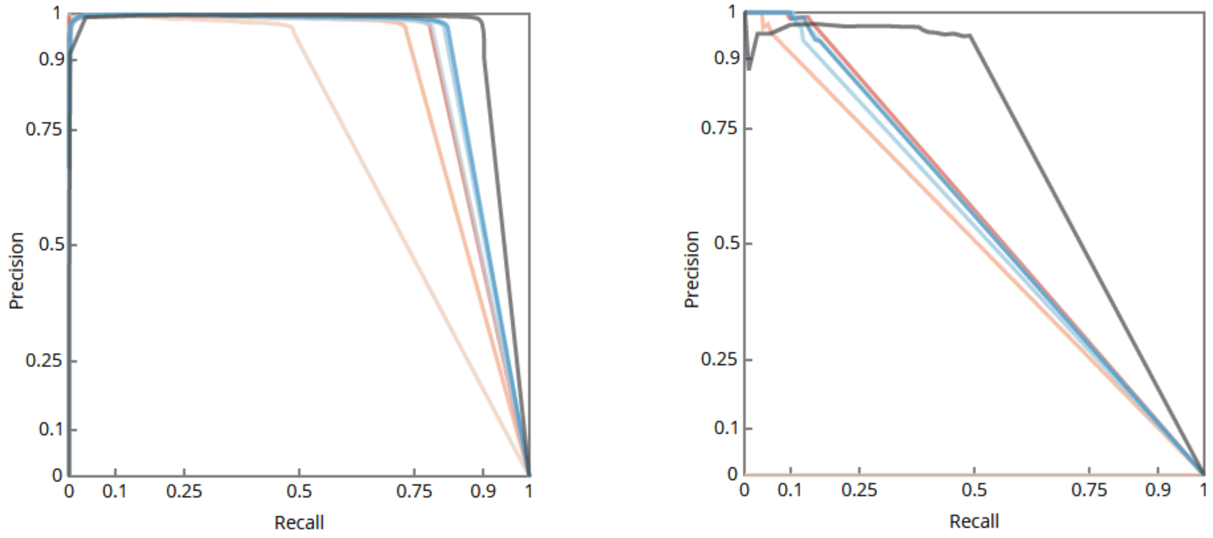

##### SRR515199

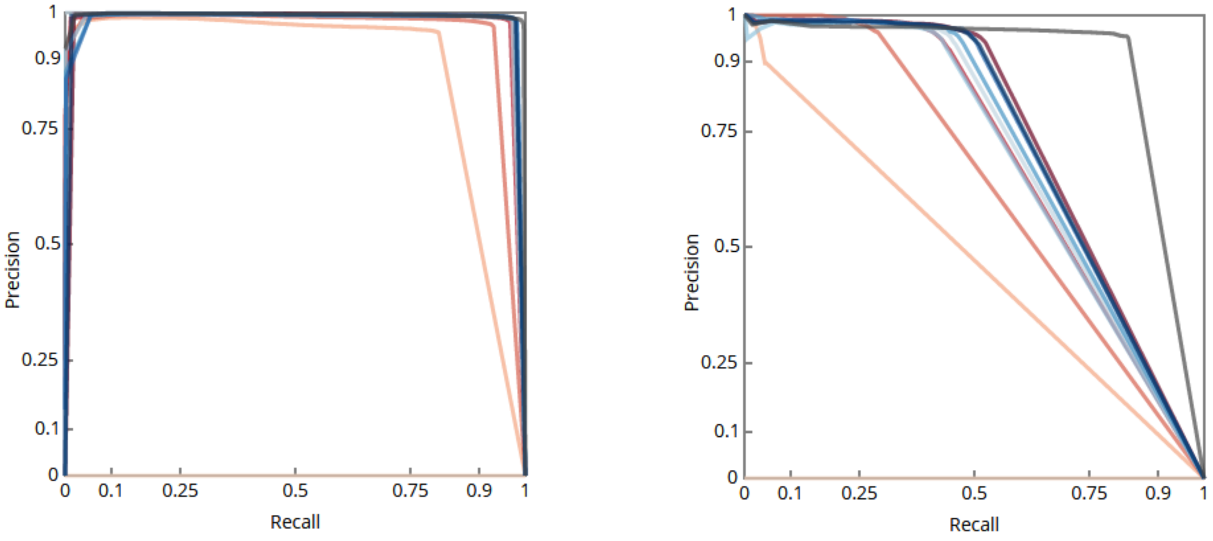

**SRR1611178**

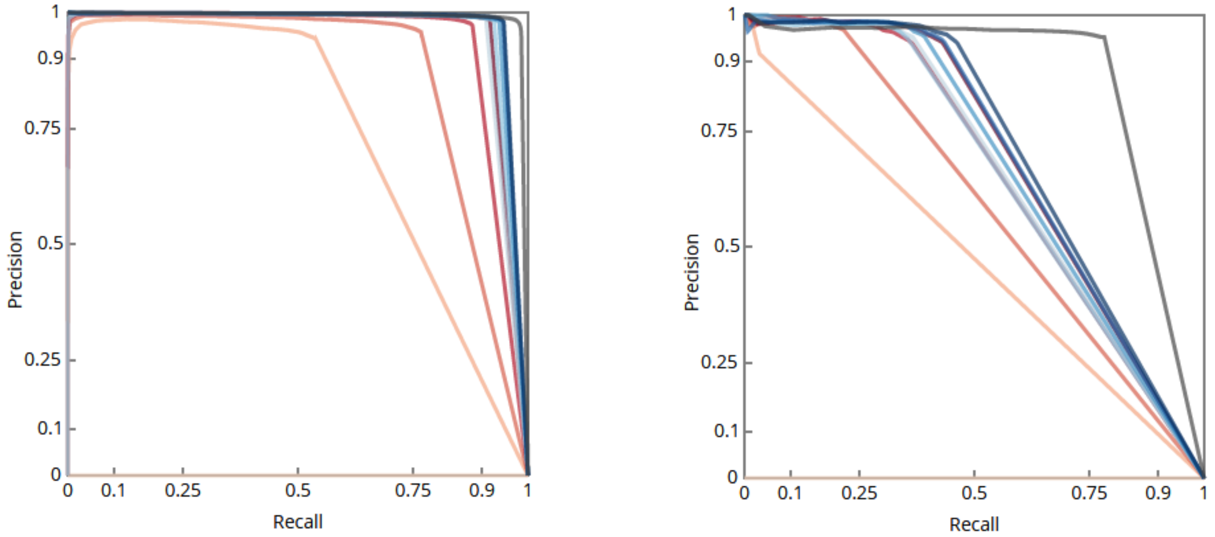

**SRR1611179**

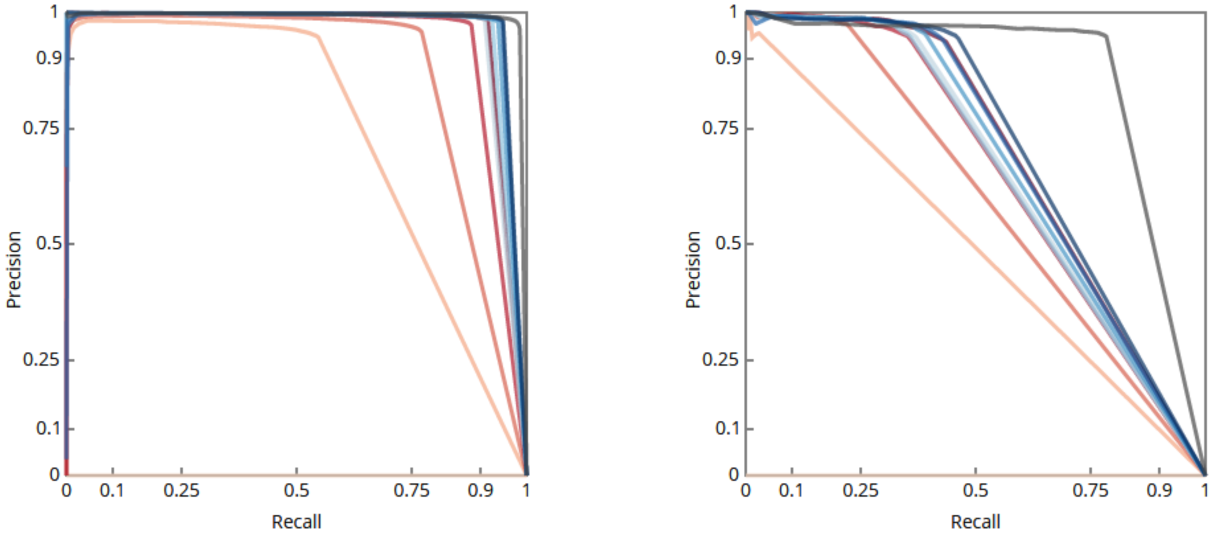

**SRR1611183**

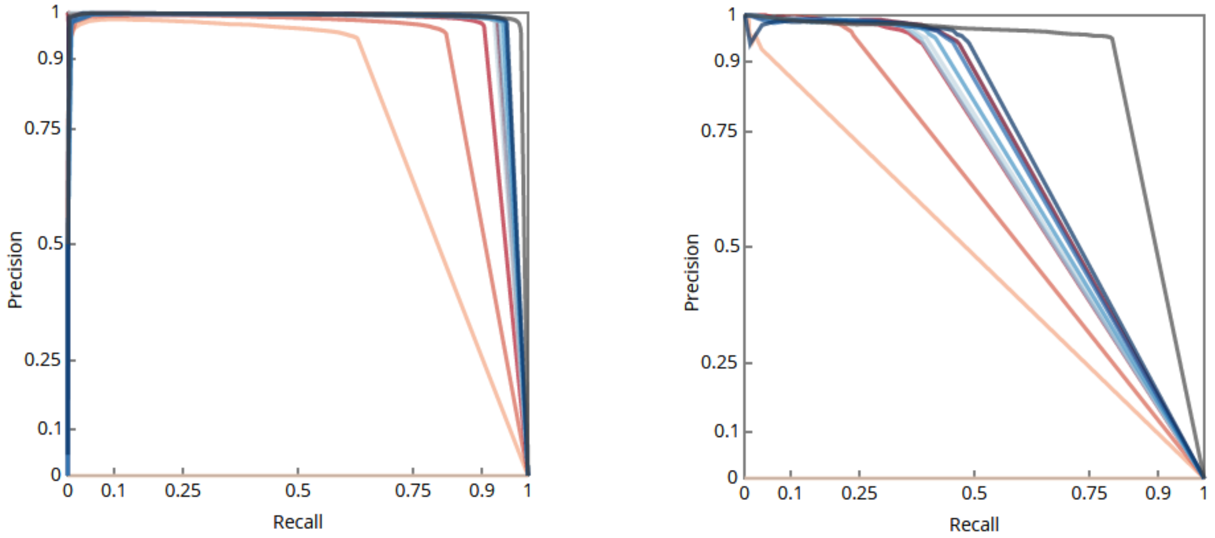

**SRR1611184**

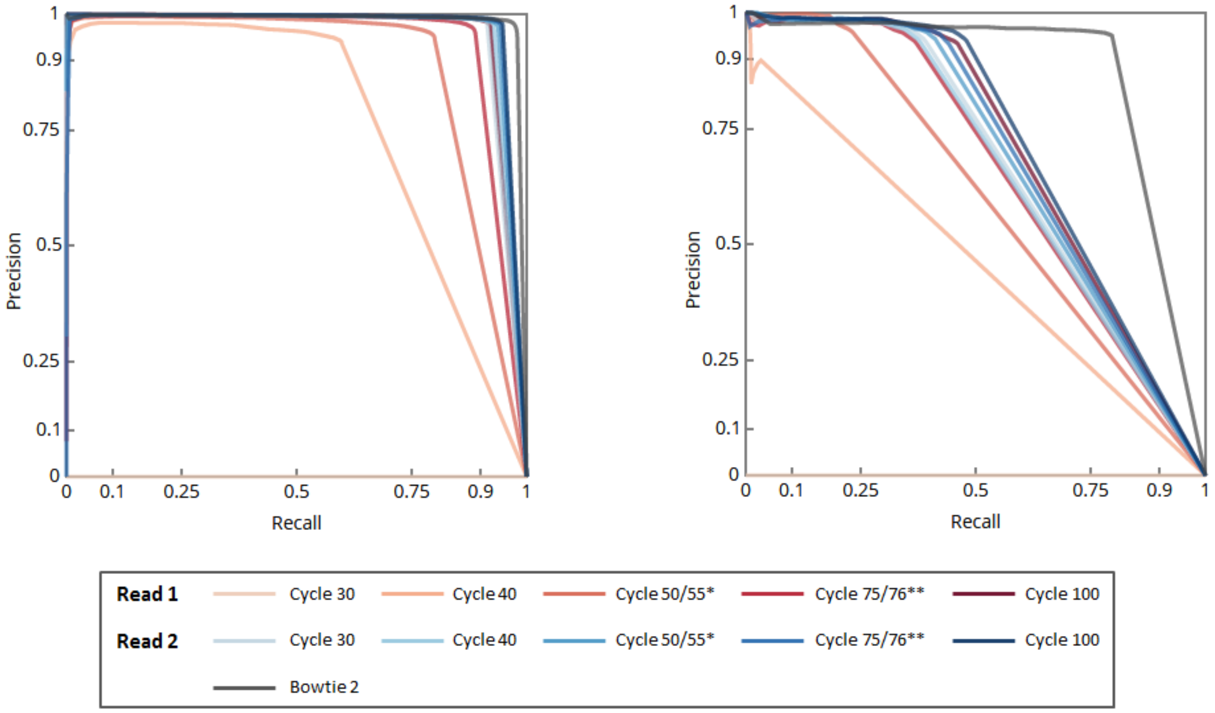

**Supplementary Fig. 1: Precision-Recall (PR) curves for seven data sets of human individual NA12878 for a real-time variant calling workflow using HiLive2 and xAtlas.** Red curves show results for the first read, blue curves show results for the second read and the gray curve show results based on read mapping with Bowtie 2. The left figure for each data set shows the PR-curve for SNPs, the right figure shows PR-curves for short indels. For SNP calling, recall increases with longer sequencing time while the precision is high from the very beginning. Final results show at least the same precision as results based on Bowtie 2 but have slightly lower recall. For short indels, an additional preprocessing-step was performed to left-align all indel positions in the read mapping results. The general tendency of the results is very similar to those for SNPs but results show much lower recall than based on Bowtie 2. This is because HiLive 2 can only find consecutive indels up to a length of 3 with the used parameter settings. As longer indels are included in the gold standard, 18-24% of gold standard indels cannot be identified based on read mapping results with HiLive 2. The particularly low recall for indels in data set SRR292250 is because of the short read length of 50bp.

#### Supplementary Fig. 2

##### SRR098401

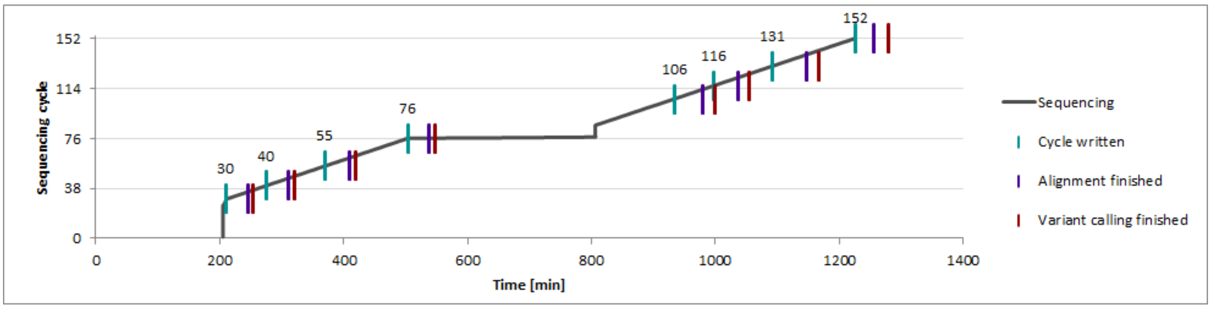

##### SRR292250

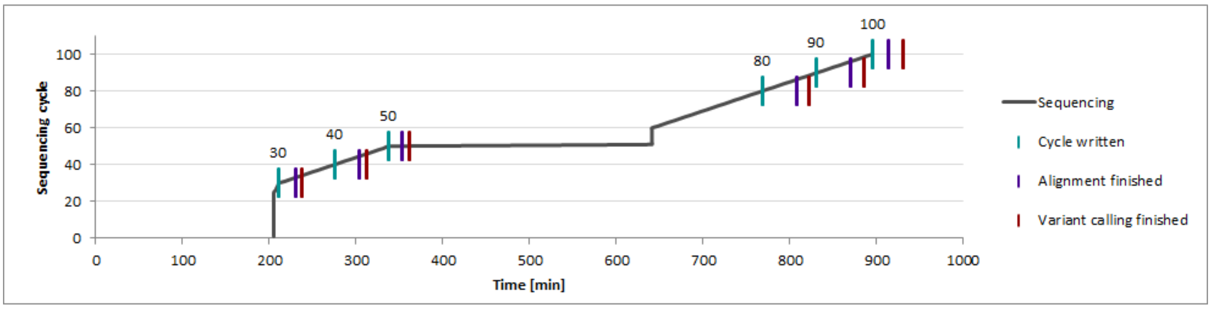

##### SRR515199

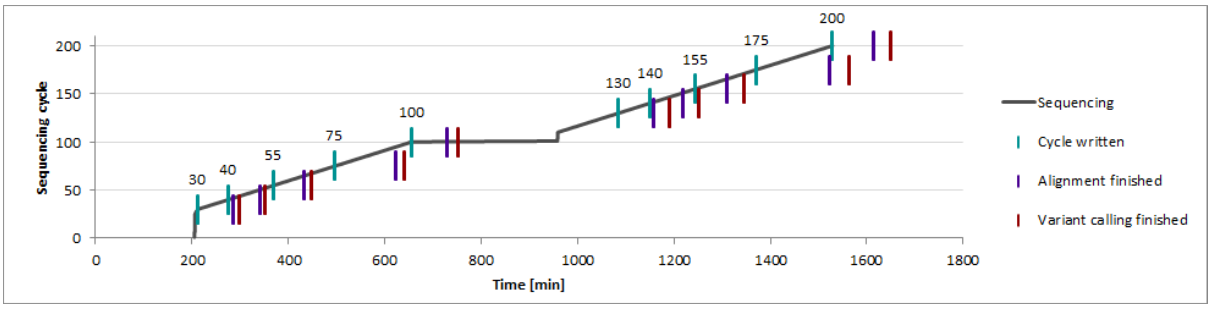

##### SRR1611178

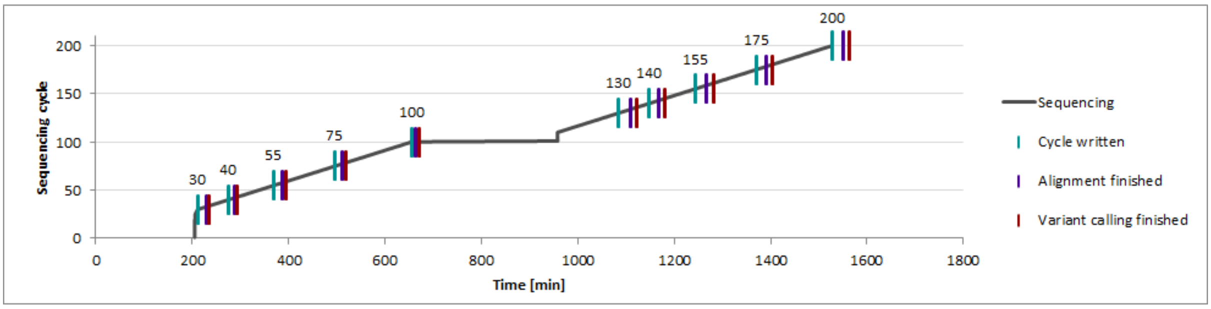

##### SRR1611179

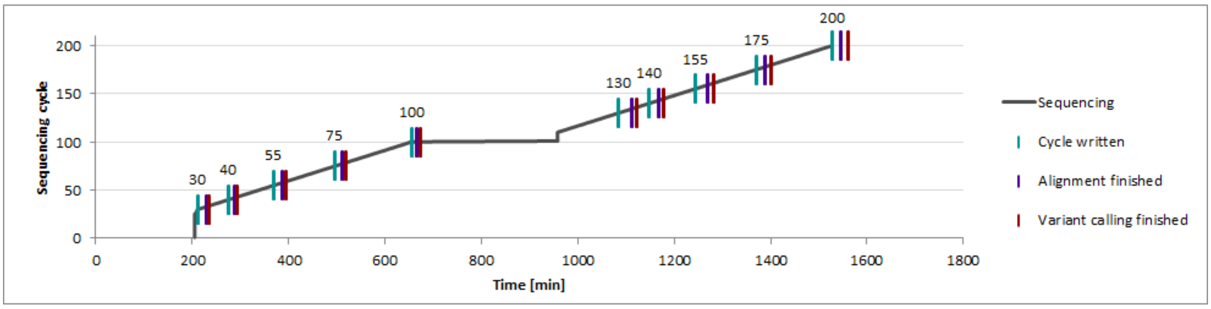

#### SRR1611183

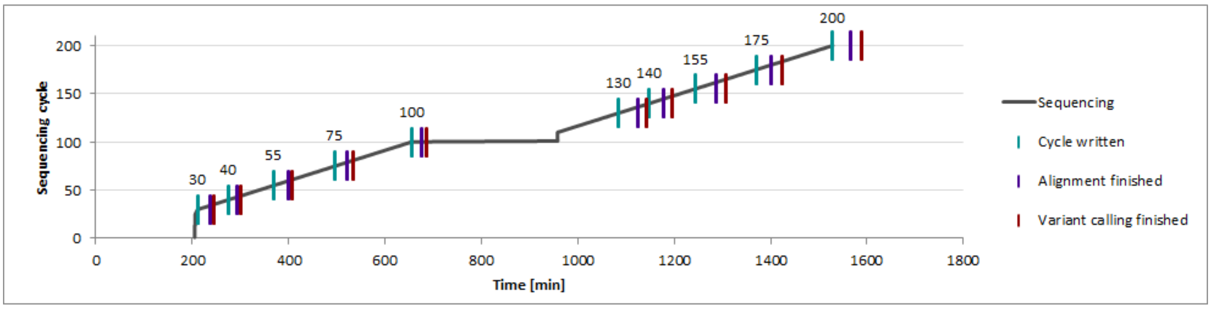

#### SRR1611184

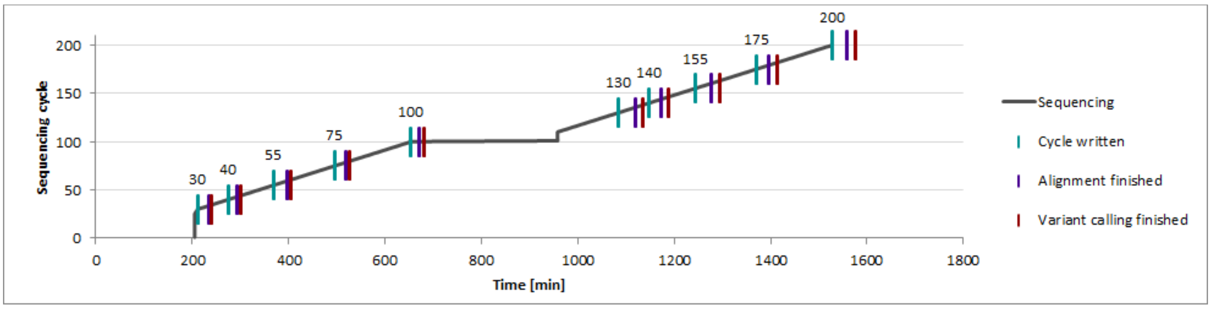

**Supplementary Fig. 2: Turnaround times for the variant calling workflow for seven data sets and different sequencing cycles.** The x-axis describes the turnaround time from starting the sequencer in minutes. The y-axis describes the sequencing cycle. The first vertical line of each data point indicates the time point when the sequencing cycle was written by the sequencing machine. The second vertical line shows when read mapping with HiLive2 finished and the third vertical line represents the availability of final variant calling results with xAtlas. The presented turnaround times do not include left-aligning indels that was necessary to call indels. Sequencing time was simulated with a simulator for illumina sequencing.

#### Supplementary Fig. 3

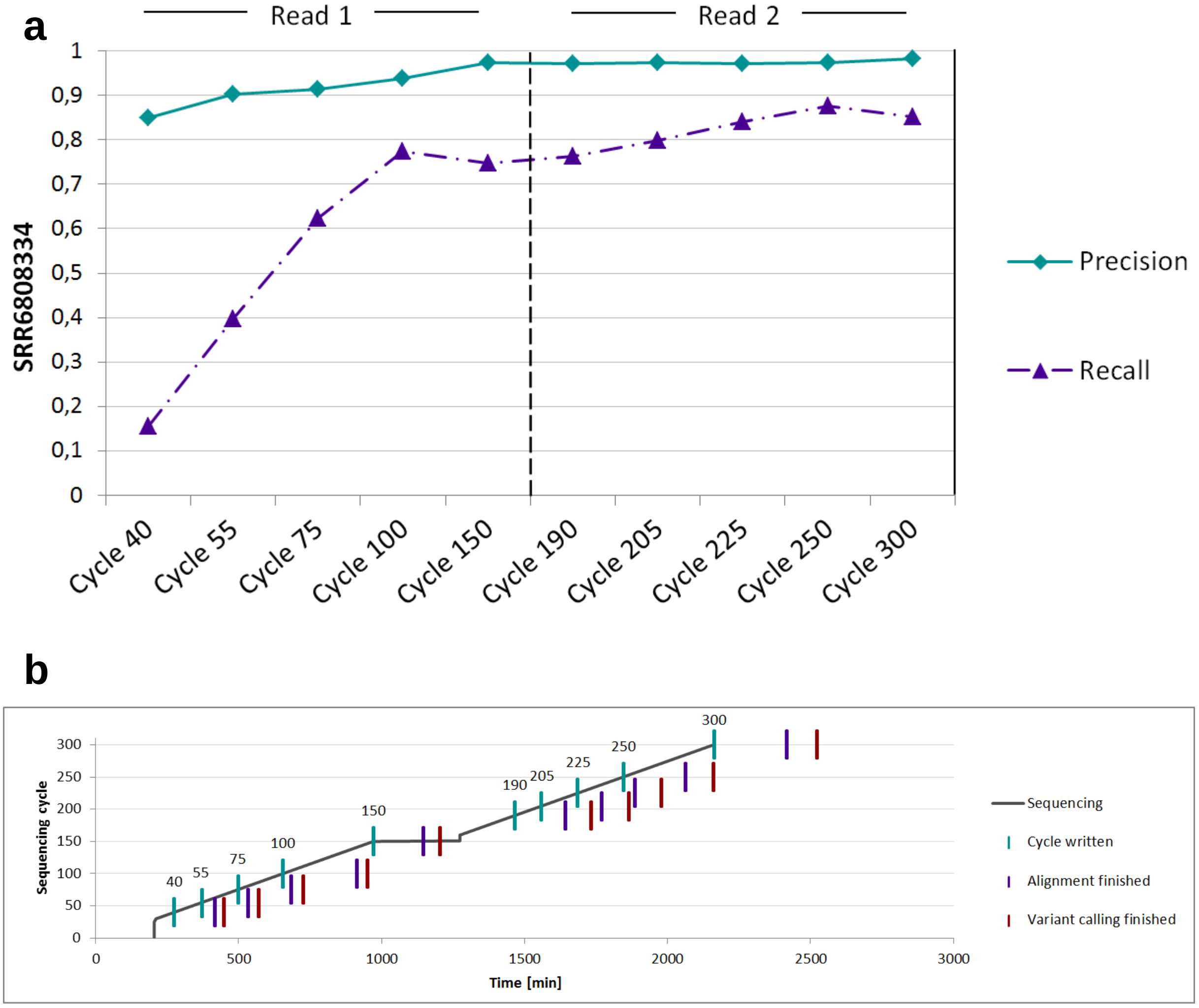

**Supplementary Fig. 3: Precision, recall (a) and turnaround times (b) for the variant calling workflow for the WGS data set SRR6808334 and different sequencing cycles. a** Precision and recall for SRR6808334 in different sequencing cycles. The x-axis denotes the output cycles. The y-axis denotes the values for precision and recall. **b** The x-axis describes the turnaround time from starting the sequencer in minutes. The y-axis describes the sequencing cycle. The first vertical line of each data point indicates the time point when the sequencing cycle was written by the sequencing machine. The second vertical line shows when read mapping with HiLive2 finished and the third vertical line represents the availability of final variant calling results with xAtlas. Sequencing time was simulated with a simulator for illumina sequencing.

### Data availability and processing

#### Human GRCh37 reference genome

##### FASTA

Obtained from NCBI via the following commands on April 3, 2018:

```
for i in $(seq 1 22) X Y; do wget --quiet ftp://ftp.ncbi.nlm.nih.gov/genomes/archive/old_genbank/Eukaryotes/vertebrates_mammals/Homo_sapiens/GRCh37/Primary_Assembly/assembled_chromosomes/FASTA/chr$i.fa.gz; done
```

```
for i in $(seq 1 22) X Y; do zcat chr$i.fa.gz >> GRCh37_all_chromosomes.fa && rm chr$i.fa.gz; done
```

```
### Shorten chromosome ids (e.g., to chr12)
awk -F '|' 'substr($0,1,1)==">"{chr=(224384769-int($2)); if ( chr==23 ) chr="X"; else if (chr==24) chr="Y"; print ">chr"chr; next}1' reference/human/GRCh37/GRCh37_all_chromosomes.fa > reference/human/GRCh37/GRCh37_all_chromosomes_shortIDs.fa
```

##### SDF

Obtained from *https://s3.amazonaws.com/rtg-datasets/references/hg19.sdf.zip* on April 18, 2018

#### SureSelect Human Exon

##### Sure Select Human Exon V2

Obtained from third party source *http://mcg.ustc.edu.cn/bsc/cnv/upload/SureSelect_Human_All_Exon_V2_Regions.tar.gz* due to unavailability from the manufacturer and processed with the following commands:

```
tar -xzf SureSelect_Human_All_Exon_V2_Regions.tar.gz
```

```
sed -i '1s/.*/track name="Target Regions" description="Agilent SureSelect DNA - HSureSelect Human All Exon V2" color=0,128,0/' SureSelect_Human_All_Exon_V2_Regions.bed
```

##### SureSelect Human Exon V4

Obtained from the manufacturer (*https://earray.chem.agilent.com/suredesign;* file *S03723314_Regions.bed*).

#### SeqCap EZ Exome

##### SeqCap EZ Exome v2.0

Obtained from the manufacturer ( *https://sequencing.roche.com/content/dam/rochesequence/worldwide/shared-designs/SeqCapEZ_Exome_v2.0_Design_Annotation_file.zip;* file *SeqCap_EZ_Exome_v2.bed*). The file contained both primary and capture targets in one file, thus we separated the primary targets using the following processing step:

```
awk 'NR>1 && $1=="track" {exit;} {print;}' SeqCap_EZ_Exome_v2.bed > SeqCap_EZ_Exome_v2_targets.bed
```

**SeqCap EZ Exome v3.0**
Obtained from the manufacturer (*https://sequencing.roche.com/content/dam/rochesequence/worldwide/resources/SeqCapEZ_Exome_v3.0_Design_Annotation_files.zip;* file *SeqCap_EZ_Exome_v3_hg19_primary_targets.bed*)

#### GIAB gold standard

##### High confidence regions

Obtained from *ftp://ftp-trace.ncbi.nih.gov/giab/ftp/release/NA12878_HG001/GIABPedigreev0.2/union13callableMQonlymerged_addcert_nouncert_excludesimplerep_excludesegdups_excludedecoy_excludeRepSeqSTRs_noCNVs_v2.19_2mindatasets_5minYesNoRatio_AddRTGPlatGenConf_filtNISTclustergt9_RemNISTfilt_RemPartComp_RemRep_RemPartComp_v0.2.bed.gz* and processed with the following commands:

```
gzip –d ftp://ftp-trace.ncbi.nih.gov/giab/ftp/release/NA12878_HG001/
GIABPedigreev0.2/union13callableMQonlymerged_addcert_nouncert_
excludesimplerep_excludesegdups_excludedecoy_excludeRepSeqSTRs_noCNVs_
v2.19_2mindatasets_5minYesNoRatio_AddRTGPlatGenConf_filtNISTclustergt9_
RemNISTfilt_RemPartComp_RemRep_RemPartComp_v0.2.bed.gz

### Modify chromosome IDs to match the short ID format (e.g., chr12)
sed -e 's/^/chr/' ftp://ftp-trace.ncbi.nih.gov/giab/ftp/release/
NA12878_HG001/GIABPedigreev0.2/union13callableMQonlymerged_addcert_
nouncert_excludesimplerep_excludesegdups_excludedecoy_excludeRepSeqSTRs_
noCNVs_v2.19_2mindatasets_5minYesNoRatio_AddRTGPlatGenConf_filtNISTcluste
rgt9_RemNISTfilt_RemPartComp_RemRep_RemPartComp_v0.2.bed > hg19_high_
confidence_regions_fullchrid.bed
```

##### High confidence variants

Obtained from *ftp://ftp-trace.ncbi.nih.gov/giab/ftp/release/NA12878_HG001/GIABPedigreev0.2/NIST_RTG_PlatGen_merged_highconfidence_v0.2_Allannotate.vcf.gz* and processed with the following commands:

```
gzip -d NIST_RTG_PlatGen_merged_highconfidence_v0.2_Allannotate.vcf.gz

### Convert complex variants to primitives
vcfallelicprimitives NIST_RTG_PlatGen_merged_highconfidence_v0.2_
Allannotate.vcf > NIST_RTG_PlatGen_merged_highconfidence_v0.2_
Allannotate_allelicprimitives.vcf

### Modify chromosome IDs to match the short ID format (e.g., chr12)
grep "^#" NIST_RTG_PlatGen_merged_highconfidence_v0.2_Allannotate_
allelicprimitives.vcf > NIST_RTG_PlatGen_merged_highconfidence_v0.2_
Allannotate_allelicprimitives_renamedchromosomes.vcf

grep -v "^#" NIST_RTG_PlatGen_merged_highconfidence_v0.2_Allannotate_
allelicprimitives.vcf | sed -e 's/^/chr/' >> NIST_RTG_PlatGen_merged_
highconfidence_v0.2_Allannotate_allelicprimitives_renamedchromosomes.vcf

### Split SNPs and Indels in separated files
awk 'substr($0,1,1)=="#"{print; next;}length($4)==1 && length($5)==1{
print; next; }' NIST_RTG_PlatGen_merged_highconfidence_v0.2_Allannotate_
allelicprimitives_renamedchromosomes.vcf > NIST_RTG_PlatGen_merged_
highconfidence_v0.2_Allannotate_allelicprimitives_renamedchromosomes_snp.
vcf
```

```
awk 'substr($0,1,1)=="#"{print; next;}length($4)!=1 || length($5)!=1{
print; next; }' NIST_RTG_PlatGen_merged_highconfidence_v0.2_Allannotate_
allelicprimitives_renamedchromosomes.vcf > NIST_RTG_PlatGen_merged_
highconfidence_v0.2_Allannotate_allelicprimitives_renamedchromosomes_inde
l.vcf
```

**Gold standard variants in exome capture regions**

For each exome capture region file, two individualized gold standard variant files (separated SNPs and Indels) were created by executing the following commands:

```
bedtools intersect -header -a NIST_RTG_PlatGen_merged_highconfidence_
v0.2_Allannotate_allelicprimitives_renamedchromosomes_snp.vcf
-b EXOME_CAPTURE_NAME.bed > gold_standard_EXOME_CAPTURE_NAME_snp.vcf
```

```
bedtools intersect -header -a NIST_RTG_PlatGen_merged_highconfidence_
v0.2_Allannotate_allelicprimitives_renamedchromosomes_indel.vcf
-b EXOME_CAPTURE_NAME.bed > gold_standard_EXOME_CAPTURE_NAME_indel.vcf
```

##### Datasets of human individual NA12878

The WES datasets SRR292250, SRR515199, SRR098401, SRR1611178, SRR1611179, SRR1611183 and SRR1611184 as well as WGS dataset SRR6808334 were obtained in compressed FASTQ format from the EBI ftp server (*ftp://ftp.sra.ebi.ac.uk/vol1/fastq/*).

#### Supplementary Methods

##### Fastq2bcl

Base call files for all data sets were created using the `fastq2bcl.py` script. Before, both reads were concatenated to enable common conversion of both reads:

```
for sample in SRR098401 SRR1611178 SRR1611179 SRR1611183 SRR1611184
SRR292250 SRR515199; do concatenate_reads.sh -z -f ${sample}/${sample}_
1.*.gz -s ${sample}/${sample}_2.*.gz > ${sample}_concatenated.fastq;
fastq2bcl.py --max_tile 2216 -o ./${sample} ${sample}_concatenated.fastq
&& rm ${sample}_concatenated.fastq; mv ./${sample}/BaseCalls ./${sample}/
BaseCalls_orig; done
```

The WGS dataset SRR6808334 was prepared in the same way, but included the option "`-l 150`" for `concatenate_reads.sh` to extend all reads to the same read length.

##### Index building

The creation of the input FASTA file is described in the section *Data availability and processing → Human GRCh37 reference genome → FASTA*:

###### HiLive2

The human index for HiLive2 was built with the `hilive-build` executable that is delivered with the HiLive2 read mapping software.

```
hilive-build -i GRCh37_all_chromosomes_shortIDs.fa -o GRCh37
```

###### Bowtie 2

The human index for Bowtie 2 was built with the `bowtie2-build` executable that is delivered with the Bowtie 2 read mapping software:

```
bowtie2-build GRCh37_all_chromosomes_shortIDs.fa GRCh37
```

##### Simulation of Illumina sequencing

Simulation of Illumina sequencing was performed using the Illumina sequencing simulator that was implemented for this study. We used the simulation model HISEQ_2500_RAPID_DUALBC while not writing base call files for the barcode cycles as those were not present in the data. Read lengths were adapted for the different data sets:

```
### SRR098401
~/scripts/simulate_run.py -o . -r 76R,8b,8b,76R -m
HISEQ_2500_RAPID_DUALBC BaseCalls_orig/

### SRR292250
~/scripts/simulate_run.py -o . -r 50R,8b,8b,50R -m
HISEQ_2500_RAPID_DUALBC BaseCalls_orig/

### All other WES data sets
~/scripts/simulate_run.py -o . -r 100R,8b,8b,100R -m
HISEQ_2500_RAPID_DUALBC BaseCalls_orig/

### SRR6808334
~/scripts/simulate_run.py -o . -r 150R,8b,8b,150R -m
HISEQ_2500_RAPID_DUALBC BaseCalls_orig/
```

##### Read mapping with HiLive2

HiLive2 was executed using individualized configuration files specifying the read length and output cycles for each data set. Besides technical settings such as input and output directories and the number of lanes and tiles, fast mode was activated and the number of threads was set to 64 for WES datasets. For the WGS data set, parameters were adapted manually to achieve better runtime behavior and 96 tiles and threads were used:

```
### Exemplary WES configuration file for data set SRR098401
lanes=1
max-tile=2216
num-threads=64
out-dir=./out
temp-dir=./temp
bcl-dir=./BaseCalls
reads=76R,76R
out-cycles=30,40,55,76,106,116,131,152
align-mode=fast
```

```
### Configuration file for WGS data set SRR6808334
lanes=1
max-tile=2316
num-threads=96
out-dir=./out
temp-dir=./temp
bcl-dir=./BaseCalls
reads=150R,150R
out-cycles=40,55,75,100,150,190,205,225,250,300
align-mode=very-fast
min-as=-16
anchor-length=30
seeding-interval=30
error-interval=30
max-gap-length=1
max-softclip-length=50
```

HiLive2 was then executed with the following command:

```
hilive --config hilive_settings.ini
```

Turnaound time for a sequencing cycle was determined by measuring the elapsed time from starting the sequencing simulation script and the time stamp of the output file for the respective cycle.

##### Read mapping with Bowtie 2

Bowtie 2 was executed with default parameters using the FASTQ files from EBI as input:

```
bowtie2 -p 10 -x GRCh37 -1 $(realpath *_1.fastq.gz) -2 $(realpath
*_2.fastq.gz) | samtools view -bS - > bowtie2_results.bam
```

##### Variant Calling with xAtlas

Variant calling with xAtlas was performed using a specialized script `run_xatlas.sh` to optimized runtime for human data sets. Therefore, the read mapping results are separated by chromosome. Sorting and variant calling are performed for each chromosome individually in parallel. Variants are only called on the regions specified by the respective exome capture. For WGS dataset SRR6808334, no exome capture file was specified. When all threads finished, the output files of the different chromosomes are joined. Runtime was measured using bash time. The following command is exemplary for data set SRR098401 and read mapping results with HiLive2 after 30 sequencing cycles:

```
time ~/scripts/run_xatlas.sh -i hilive_out_cycle30_undetermined.bam -r
GRCh37_all_chromosomes_shortIDs.fa -s SRR098401 -p hilive_out_cycle30_
undetermined -t 24 -d hilive_out_cycle30_undetermined_xatlas_temp -x '-c
SureSelect_Human_All_Exon_V2_Regions.bed'
```

For determining turnaround time, the *real* entry of the bash time output was used (rounded to minutes).

For obtaining improved variant calling results for indels based on HiLive2 read mapping, an additional preprocessing step was performed to left-align all indels in the read mapping results. The additional parameters (red) activate `GATK LeftAlignIndels` and necessary preprocessing in the script:

```
time ~/scripts/run_xatlas.sh -i hilive_out_cycle30_undetermined.bam -r
GRCh37_all_chromosomes_shortIDs.fa -s SRR098401 -p hilive_out_cycle30_
undetermined -t 24 -d hilive_out_cycle30_undetermined_xatlas_temp -x '-c
SureSelect_Human_All_Exon_V2_Regions.bed' -l -G 'java -Xmx5G -jar
./GenomeAnalysisTK.jar' -P 'picard-tools'
```

##### Evaluation with RTG Tools

Variant call files obtained from xAtlas and gold standard files are compressed and indexed with bgzip and tabix:

```
bgzip VARIANT_FILE.vcf && tabix VARIANT_FILE.vcf.gz
```

The creation of the high confidence regions file `hg19_high_confidence_regions_fullchrid.bed` is described in *GIAB gold standard → High confidence regions*.

`rtg vcfeval` is executed with the following parameters:

```
rtg vcfeval -b GOLD_STANDARD.vcf.gz -c VARIANT_FILE.vcf.gz -o
VARIANT_FILE_vcfeval -t hg19.sdf -e hg19_high_confidence_regions_
fullchrid.bed --squash-ploidy --all-records -f QUAL -T 24
```

##### Precision-Recall curves

Precision-Recall curves were created from the RTG Tools output using the visualization library $D^3$ (Bostock et al., $D^3$: Data-Driven Documents, IEEE Trans Vis Comput Graph, 2011). For visualization, a precision value was calculated for recall values in intervals of 0.0001 by linear interpolation based on the precision and recall values obtained from RTG Tools. To obtain a curve for the complete recall span [0,1], additional values were added to the input data points. The first additional data point in format (recall, precision) was set to $(0,p_1)$, $p_1$

being the precision of the lowest measured recall. The second additional data point was (1,0). Area under a precision-recall curve was calculated by the sum of the areas in recall intervals of 0.0001 which is exact for linearly interpolated input data with the same intervals.
